# Pathogen-Host Adhesion between SARS-CoV-2 Spike Proteins from Different Variants and Human ACE2 Probed at Single-Molecule and Single-Cell Levels

**DOI:** 10.1101/2022.06.05.493249

**Authors:** Xiaoxu Zhang, Jialin Chen, Bixia Hong, Haifeng Xu, Pengfei Pei, Long Chen, Yigang Tong, Shi-Zhong Luo, Huahao Fan, Chengzhi He

## Abstract

Pathogen-Host adhesion is considered the first step of infection for many pathogens such as bacteria and virus. The binding of the receptor binding domain (RBD) of SARS-CoV-2 Spike protein (S protein) onto human angiotensin-converting enzyme 2 (ACE2) is considered as the first step for the SARS-CoV-2 to adhere onto the host cells during the infection. Within three years, a number of variants of severe acute respiratory syndrome coronavirus 2 (SARS-CoV-2) have been found all around the world. Here, we investigated the adhesion of S Proteins from different variants and ACE2 using atomic force microscopy (AFM)-based single-molecule force spectroscopy (SMFS) and single-cell force spectroscopy (SCFS). We found that the unbinding force and binding probability of the S protein from Delta variant to the ACE2 was the highest among the variants tested in our study at both single-molecule and single-cell levels. Molecular dynamics simulation showed that ACE2-RBD (Omicron) complex is destabilized by the E484A and Y505H mutations and stabilized by S477N and N501Y mutations, when compared with Delta variant. In addition, a neutralizing antibody, produced by immunization with wild type RBD of S protein, could effectively inhibit the binding of S proteins from wild type, Delta and Omicron variants onto ACE2. Our results provide new insight for the molecular mechanism of the adhesive interactions between S protein and ACE2 and suggest that effective monoclonal antibody can be prepared using wild type S protein against the different variants.

## Introduction

Severe acute respiratory syndrome coronavirus 2 (SARS-CoV-2) has rapidly spread to the entire world and become a devastating pandemic^1,2^. As the SARS-CoV-2 is an enveloped, positive-stranded RNA virus, mutation with increased infectivity and transmission is likely to happen during the evolution.^3,4^ SARS-CoV-2 infects human host cells by an initial adhesive interaction of its receptor-binding domain (RBD) located in the C-terminal of the S1 subunit of spike protein (S protein) and angiotensin converting enzyme 2 (ACE2), the receptor on human cells, as shown in Figure 1A and B^2,5^. The mutations in RBD of S protein may result in higher adhesion force and thus affect infectivity and virulence. Recently, a number of mutations were identified on the S protein in the SARS-CoV-2 variants, which have been reported with different infectivity and pathogenicity.^1^ In particular, an N501Y mutation in the RBD of S protein involve in Alpha (B.1.1.7), Beta (B.1.351) and Gamma (P1) has an increased affinity and higher virulence to ACE2. ^4,6-8^ Meanwhile, the K417N/T and E484K mutations in the RBD of Beta (B.1.351) and Gamma (P1) may result in conformational changes.^1^ The Delta variant (B.1.617.2) harbors L452R and T478K mutations in the RBD, which could enhance the stability of S protein and the affinity to ACE2.^1,9,10^ Compared with Delta variant, Omicron variant (BA.1) contains more mutations, among which there are 10 residues located on the interface of the S protein and ACE2. In the past two years, the Delta and Omicron variants have become two of the most popular variants among all the variants of concern (VOCs). Currently, it is believed that the Omicron variant is more infectious but less pathogenic than Delta variant.

**Figure 1.**
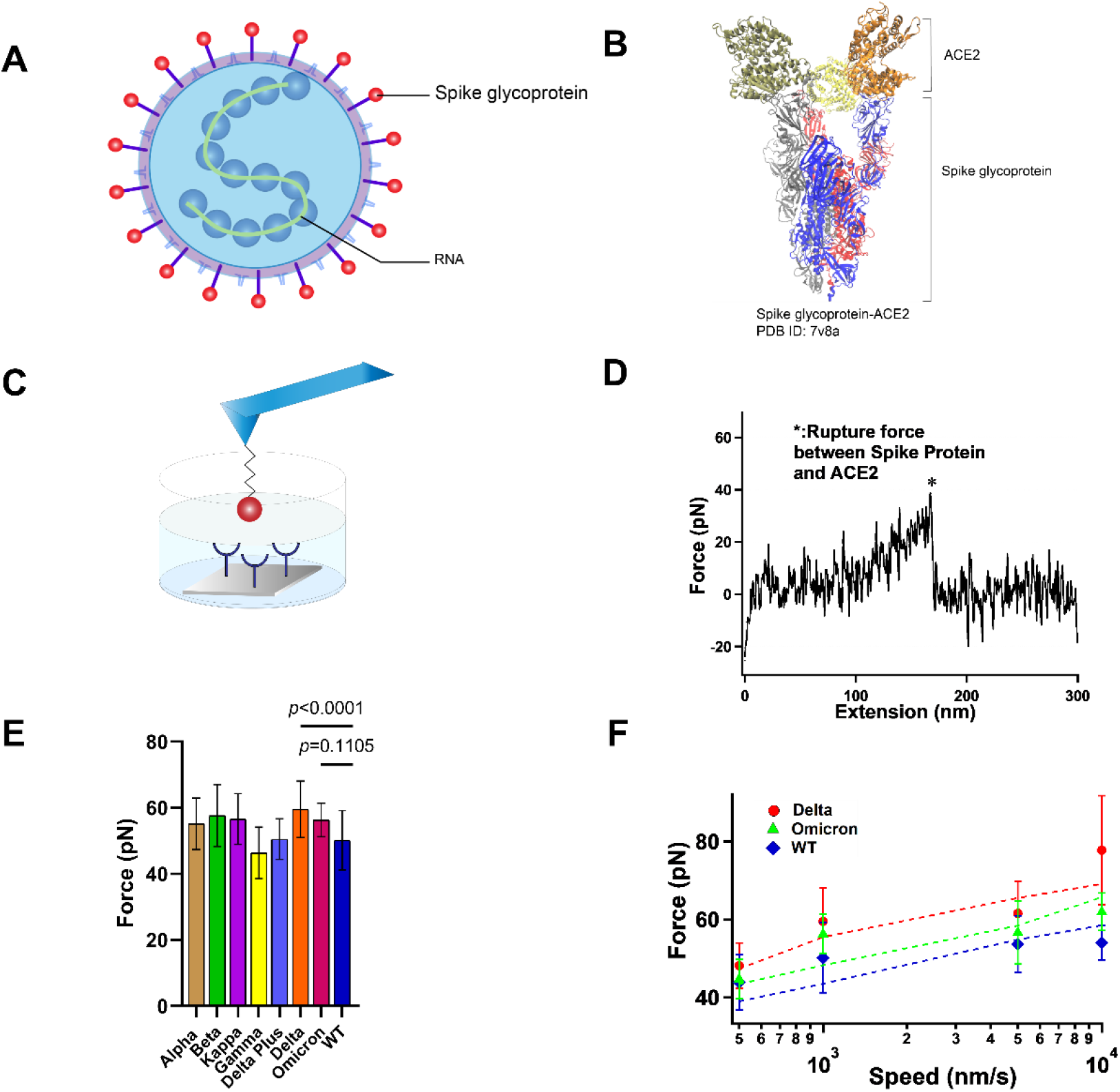
The interaction between S protein and ACE2 was studied by SMFS. (**A**). Schematic diagram of SARS-COV-2 particle, an enveloped ssRNA virus expressing spike glycoprotein (S) on its surface. (**B**). Crystal structure of Delta variant (B.1.617.2) in complex with three ACE2. (**C**). Schematic of measurement of adhesive interaction between S protein and ACE2 using SMFS. The purified S protein and ACE2 were attached to probe and substrate, respectively. (**D**). A representative force-extension curve showing specific adhesive event labeled by star. (**E**). The adhesive forces between S proteins from different variants and ACE2. (**F**) Pulling speed dependency of adhesive forces and Monte Carlo simulations for extracting the kinetic parameter. Solid dots represent experimental data and dotted lines represent simulated results.

The interaction between of SARS-CoV-2 S protein and ACE2 has been studied using various methods such as Cryo-EM, X-ray, surface plasmon resonance (SPR), flow cytometry and single-molecule force spectroscopy.^2,4,11-16^ Here we investigated the adhesive interactions between S protein from different variants and ACE2 using atomic force microscopy-based single molecular force spectroscopy (SMFS) and Fluid microscopy-based single cellular force spectroscopy (SCFS). Molecular dynamics simulation was also used to investigate the binding mechanism for ACE2 and RBD from Delta and Omicron variants. In addition, we evaluated the neutralization efficiency of a monoclonal antibody (mAb) obtained by immunization of mouse with the wild type (WT) RBD. We observed that the addition of mAb resulted in significant reduction of binding onto ACE2 for the wild type, Delta and Omicron variants.

## Results and Discussions

### Single-Molecule Adhesive Interactions between S proteins and ACE2

Single-molecule force spectroscopy (SMFS) has become a powerful tool to characterize the mechanical property of macromolecules^17-20^ and the protein-protein interactions^2,4,13,14,21^ at single-molecule level. Here we use atomic force microscopy (AFM) based SMFS to measure the mechanical unbinding force between ACE2 and S proteins from eight different variants. We covalently attached purified ACE2 and S proteins to AFM probe and glass substrate, respectively, using a 20 kDa NHS-PEG-NHS linker (Figure 1C). The probe was approaching to and retracting from the substate at a constant speed of 1 um/s in PBS buffer and at room temperature. A typical relationship between force and extension is shown in Figure 1D. The force-extension curves (FECs) were fitted with the worm-like chain (WLC) model using the persistence length of 0.4 nm for the PEG linker. The single force-rupture events correspond to the mechanical unbinding of S protein and ACE2. Representative histograms of rupture force distribution for the interaction between ACE2 and different S proteins are shown in Figure S1 and the mutations of different variants are listed in Table S1. The unbinding forces of ACE2 and S proteins are summarized in Figure 1E and all the unbinding forces are in the range from ∼40 to 60 pN, which is in agreement with previous studies^2,4,13,14^. S proteins from all variants except the Gamma variant had greater unbinding forces to ACE2 than the wild type, although the difference between different variants is small. Among these variants, the Delta variant has the highest unbinding force to ACE2, which can reach up to 60±9 pN (n=157).

To extract the kinetics of the unbinding process, we conducted the dynamic force spectroscopy experiments at various pulling rates on S proteins from wild type, Delta and Omicron variants (Figure 1F). According to the Bell-Evans model and Monte Carlo simulation^22-24^, the dissociation rate constant *k*_0_ in the absence of a pulling force and the distance from native state to transition state *Δx*_β_ were estimated as shown in Table 1. Our results showed that all three S proteins has similar *Δx*_β_ and dissociation rate constant, suggesting that they have similar unbinding pathway and mechanical stability, although WT S protein - ACE is the least stable. The stronger adhesion between the Delta variant and the ACE2 may contribute to the stronger cell invasion ability of the Delta variant compared to the other variants.

**Table 1.** Kinetics extracted from dynamic force spectroscopy experiments

| S Protein-ACE2 | $k_0$ ($s^{-1}$) | $\Delta x_\beta$ (nm) |
| --- | --- | --- |
| WT S Protein-ACE2 | 0.075 | 0.71 |
| Delta S protein-ACE2 | 0.026 | 0.68 |
| Omicron S protein-ACE2 | 0.029 | 0.73 |

### Single-Cell Adhesion between S Proteins and Cells

Force spectroscopy can not only be used at single-molecule level, but it can be expanded to single-cell level as well.^2,14,25^ Atomic force microscopy is a useful tool to conduce single-cell force spectroscopy (SCFS) to characterize the interaction between a single cell and a surface as it has a broad range of measurable forces from pN to nN.^26 27,28^ SCFS has previously been used in a variety of applications, such as studying the interactions between cells and substrate materials or hydrogels and mechanisms of adhesion between cells and extracellular matrix.^26,29,30^ To further study the interaction between S proteins from different variants and ACE2 at cell level, purified S proteins and Vero cells with high expression of ACE2 were studied using SCFS. HEK293T cells were used as control cells because ACE2 protein is less expressed on the HEK293T cell surface. In addition, BSA coated and blank glass substrates were chosen as control surfaces for cell adhesion.

In our experimental setup, the attachment of a single cell to the tip of cantilever was achieved by added a negative pressure of 50 mbar from a pump, and the probe is a micropipette with an aperture of 4um. The S proteins were immobilized on glass substrate using 20 kDa NHS-PEG-NHS linker the same way as the SMFS experiments. The cell-attached probe approached the substrate at a rate of 2 um/s and contacted with the substrate for 5 s allowing the ACE2 on the cells to establish stable adhesion to S protein coated substrate, then retracted at same rate to 30um (Figure 2A). When the cell is retracted from the S proteins coated surface, the adhesion force is precisely measured. A typical force-extension curves (FECs) were shown in Figure 2B and their maximum adhesion force was recorded. The rupture events occurred at the highest force corresponds to the detachment of the bulk of cell from substrate.

**Figure 2.**
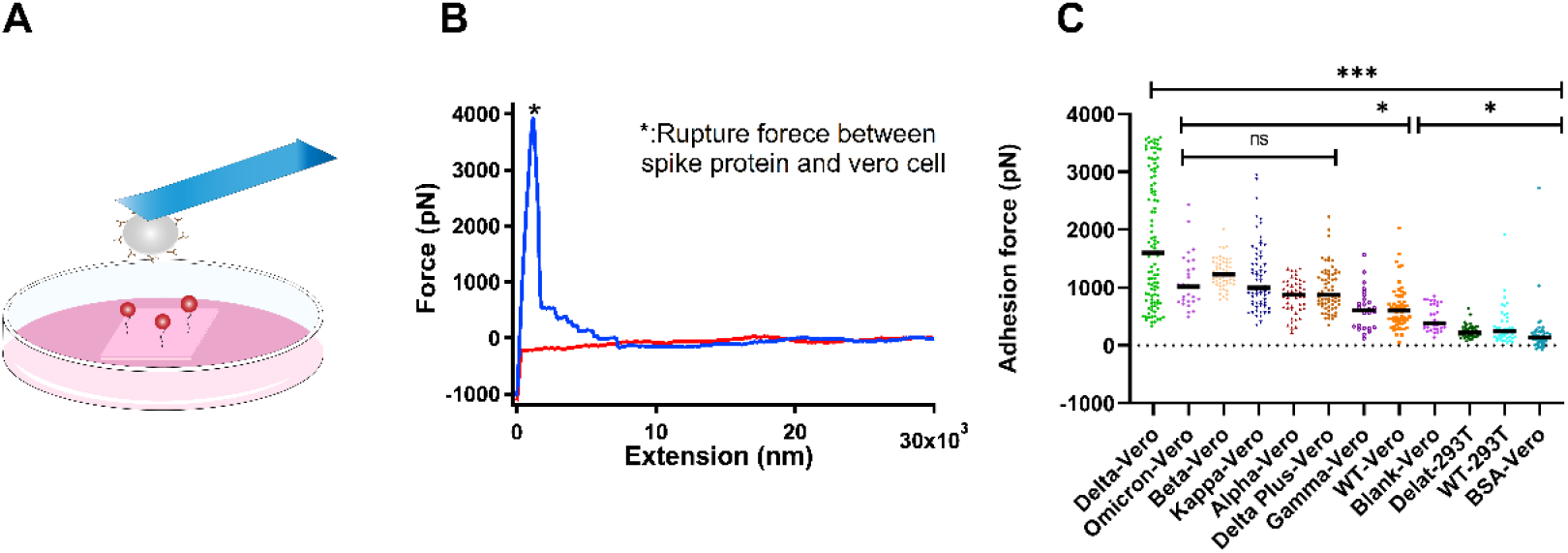
Experimental results of SCFS. (**A)**. Schematic of SCFS experiments. (**B)**. A representative force-extension curve of the detachment of Vero cell from S protein coated surface. (**C)**. Adhesion force between cells and S proteins from different variants.

The adhesion forces between cells and different surfaces were summarized in Figure 2C. The S proteins from all the variants including wild type showed higher adhesion forces than the control groups, indicating that the S proteins indeed interact with ACE2, which is consistent with the results from previous studies.^2,31^ Moreover, the results of SCFS is in agreement with our SMFS. The Delta variant showed the highest adhesion force among all the variants. No significant differences in the adhesion force were found for the Omicron variant compared with most other variants, although it is higher than wild type. Due to the multiple interactions of ACE2 on cells and the S proteins on the substrate, the adhesion force quantified by SCFS (nN level) is much higher than the single-molecule unbinding force of S protein and ACE2 (pN level).

### Inhibitory effect of neutralizing antibody on S proteins

The SARS-CoV-2 invasion of human host cells is mediated by an interaction between the receptor binding domain on the S1 subunit of the glycoprotein anchored on the virus surface and human angiotensin-converting enzyme (hACE2).^4^ The potent neutralizing antibodies (NAbs), directed to the RBD, has been administered prophylactically after exposure to infectious virus, and the mechanism of neutralization is the competitive binding of NAbs and S proteins to receptors.^32^ Various recombinant monoclonal antibodies (mAbs) are being tested in therapy, with targets to some of the responses caused by SARS-CoV-2.^33,34^ Here, we tested the binding inhibition effect of IgG1 mAb obtained from a mouse immunized with recombinant WT SARS-CoV-2 S protein using SMFS. The glass substrate was coated with ACE2 using 20 kDa PEG linker. The AFM probes modified with S protein was incubated with mAb and then approached to substrate at a constant speed of 1 um/s. When the tip contacted with the substrate for 100 ms to react with ACE2 on the surface, it was retracted at the same velocity (Figure 3A). The experiments without addition of mAb were designed as control. The force-extension curve was shown in Figure 3B. We evaluated the neutralization effect of the antibody by measuring the binding probability (BP) of S protein and ACE2 before and after adding the antibody. We observed a significant reduction of the BP when mAb was added (Figure 3C). In addition, we studied the inhibition effect of the antibody at single-cell level by measuring the BP between different S proteins and Vero cells with high expression of ACE2 receptor (Figure 3D). The representative FEC for the binding of S protein and Vero cell was shown in Figure 3E. After addition with mAb, lower BP was observed (Figure 3F), which is in agreement with our previous observation at single-molecule level. The BP of S protein and ACE2 coated substrate is higher than that of S protein and Vero cells. It is possible that the binding event of cell surface and receptor detected by the probe might be interfered by other components on the cell surface. Alsteens’ group have reported that ACE2-derived peptides, 9-O-Acetylated-sialic acid and neutralizing antibodies can have similar inhibitory effect on WT, alpha, beta, gamma and kappa variants of Sars-Cov-2.^2,13,14^ Here our results show that mAb can effectively inhibit the binding of S proteins from Delta and Omicron variants to ACE2.

**Figure 3.**
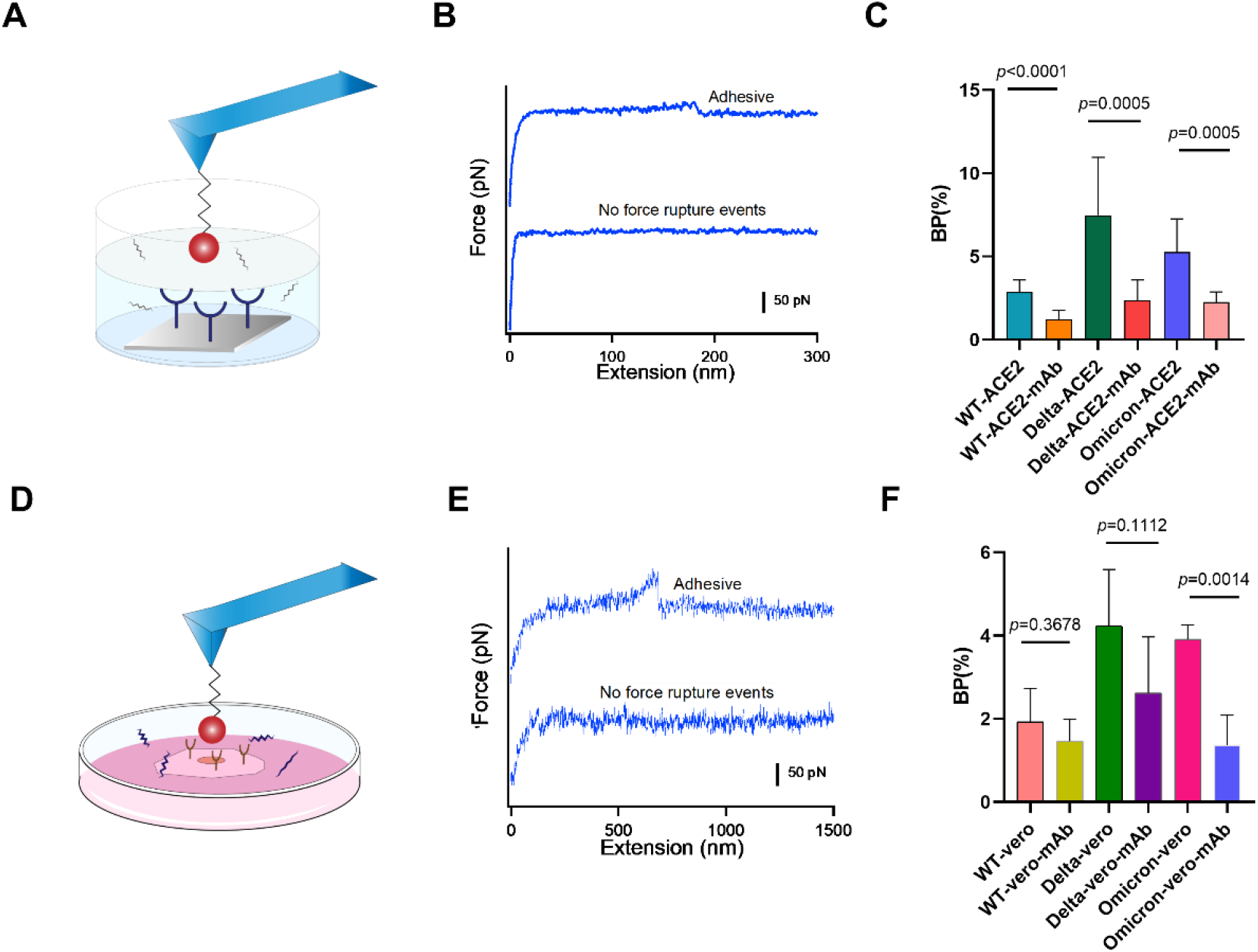
Anti-binding effects of mAb on S protein binding. **A**. Efficiency of mAb is evaluated by measuring the BP of the interaction between the S proteins and ACE2 on model surface before and after incubation of the mAb. **B**. Force-extension cures showing either nonadhesive and specific adhesive curves on model surface. **C**. Histograms of binding probabilities (BP) between purified ACE2 and S proteins (WT, Delta and Omicron). **D**. Efficiency of mAb is evaluated at the cellular level. **E**. Force-extension cures measured on Vero cells. **F**. Histograms of binding probabilities (BP) between Vero cells and S proteins (WT, Delta and Omicron).

### Binding Mechanisms of RBD-ACE2 Revealed by Molecular Dynamics Simulations

Both SMFS and SCFS experiments showed that S protein from Delta variant is the most adhesive to ACE2 protein, while the molecular mechanism is still unclear. Here we use molecular dynamics (MD) simulation to study the molecular mechanism of interaction between ACE2 and RBD from Delta and Omicron variants. Our MD simulation results showed that ACE2 and RBD from Delta and Omicron variants have similar interaction energy (Figure S2). The mutations on the RBD at the ACE2-RBD interface were summarized in Table S1. We then analyzed the interactions between ACE2 and some key residues (different in Delta and Omicron variants) on RBD at the ACE2-RBD interface as shown in Figure 4. As shown in Figure 4C, we found that K31-E484 and E37-Y505 in the ACE2-RBD (Delta) complex have strong electrostatic interactions (salt bridge and hydrogen bond, respectively), which are absent in ACE2-RBD (Omicron) complex due to E484A and Y505H mutations, while N501Y and S477N mutations stabilize ACE2-RBD (Omicron) complex by forming the π–π stacking interaction Y41-Y501 and hydrogen bond S19-N477, respectively.

**Figure 4.**
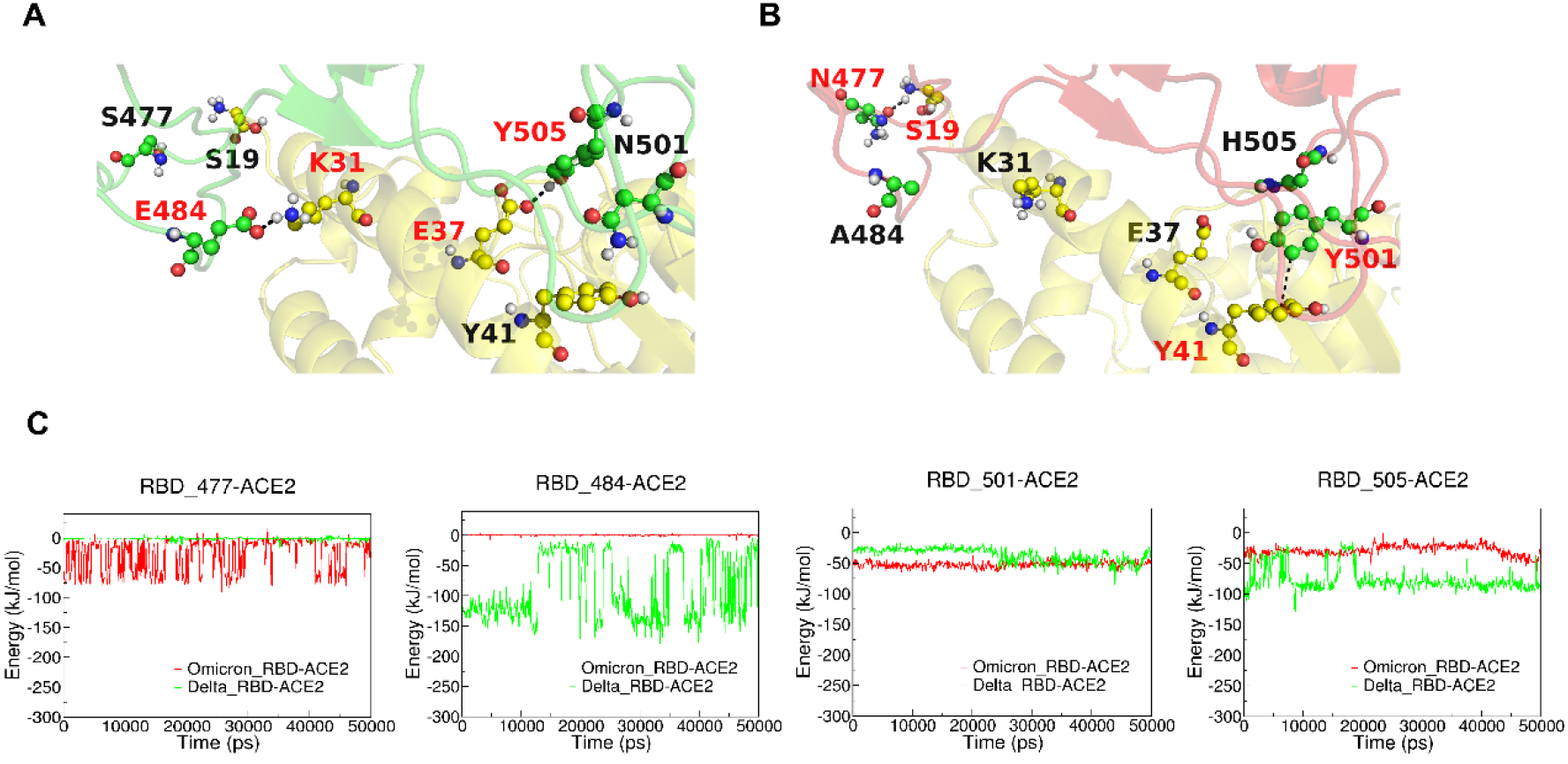
MD simulations of ACE2-RBD complex. **A** and **B**: The interactive interface of ACE2 (yellow) and RBD from Delta (**A**, green) and Omicron (**B**, red) variants. **C**: Interaction Energy of ACE2 and the individuals of 477, 484, 501 and 505 amino acid residues on RBD from Delta (green) and Omicron (red) Variants.

To mimic the force spectroscopy experiments, we performed steered molecular dynamics (SMD) simulations by pulling the RBD away from the ACE2-RBD (Delta and Omicron variants) complexes. Figure 5A shows the process from the binding state of RBD and ACE2 to the dissociation state and the dissociation occurs in 5 ns. The highest pulling force occurred at ∼3000 ps and the extension of ∼1 nm as shown in Figure 5B. Our SMD results showed that the ACE2-RBD (Delta) has slightly higher unbinding force (Figure 5C), which is consistent with our SMFS experiments. Interactions between ACE2 and key residues on RBD were analyzed as shown in Figure 5D. Similar to our MD results, in the SMD simulations, E484-ACE2 and Y505-ACE2 in the ACE2-RBD (Delta) are more stable upon pulling than A484-ACE2 and H505-ACE2 in ACE2-RBD (Omicron), while the N477-ACE2 and Y501-ACE2 in ACE2-RBD (Omicron) are more stable than S477-ACE2 and N501-ACE2 in ACE2-RBD (Delta). Zheng et al reported that N501Y mutation results in higher binding affinity of S protein from Beta variant to ACE2, which is in agreement with our results here.

**Figure 5.**
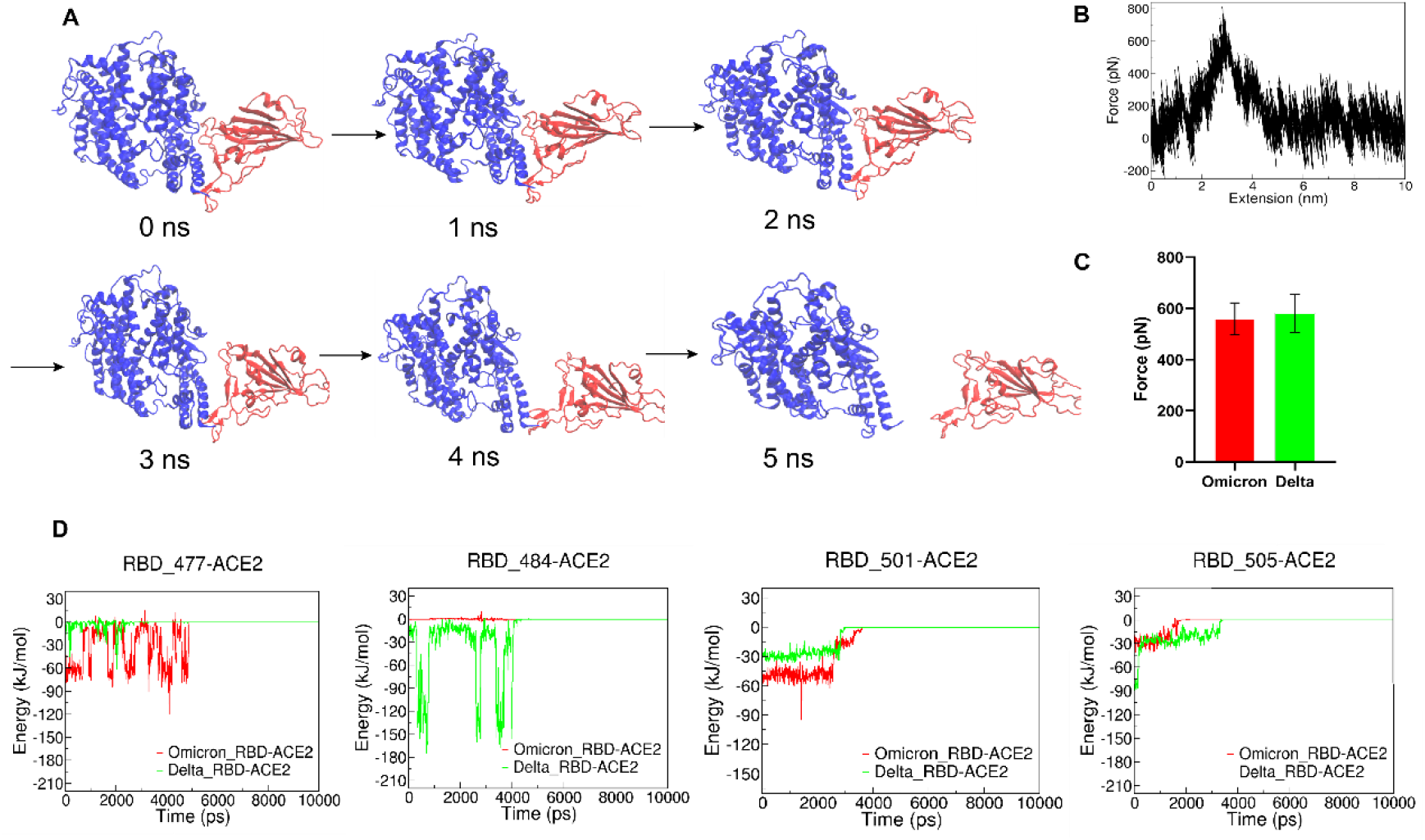
SMD simulation of RBD and ACE complex. **A**. Schematics of pulling RBD away from ACE2. **B**. Representative force-extension curve of pulling RBD away from ACE2. **C**. Unbinding forces of ACE2 and RBD from Delta and Omicron variants. **D**. Interaction energy between ACE2 and individual amino acid residues on RBD.

### Conclusions

Our results suggest S protein from Delta variant has stronger adhesive interaction than S proteins from other variants. Omicron variant the adhesion force was not higher and the mutated residues on RBD did not result in stronger interaction with ACE2 when compared with Delta variant. Mutations on RBD from Omicron variant have complex effect on the binding affinity to ACE2. E484A and Y505H mutations destabilize the binding, while N501Y and S477N stabilize the binding. It is possible that this adhesion behavior is related to the stronger infectivity and lower virulence of Omicron variant. In addition, we found that the monoclonal antibody produced using WT Sars-Cov-2 can effectively inhibit the binding of S proteins from Delta and Omicron to ACE2 at both single-molecule and single-cell levels. Our result suggests that although vaccines produced using wild type S protein has showed much lower efficiency to the omicron variant in reality, effective monoclonal antibody can be prepared using wild type S protein against the Delta and Omicron variants by inhibiting the pathogen-host adhesion.

## Methods

### Proteins

All the SARS-CoV-2 S proteins (Omicron: SPN-C52Z, Delta: SPN-C82Ec, Delta Plus: SPN-C52Ht, Alpha: SPN-C82E5, Beta: SPN-C82E4, Kappa: SPN-C82E7, Gamma: SPN-C82E6 and Wild Type: SPN-C82E9), ACE2 protein (AC2-H52H4) and anti-SARS-CoV-2 Spike RBD neutralizing antibody, Chimeric mAb (S1N-M122), recombinantly expressed from HEK293 cells, were purchased from ACRObiosystems.

### Cell culture

Vero cells with overexpression of ACE2 receptor and HEK293T cells without expression of receptor ACE2 were cultured in Dulbecco modified Eagle medium (DMEM), supplemented with 10% (vol/vol) fetal bovine serum (FBS), 100 U/ml penicillin and 100 ug/ml streptomycin in a humidified atmosphere with 5% CO_2_ at 37°C. Cells were cultured in T25 cell culture flasks and grown to 80-90% for force spectroscopy experiments.

### Functionalization of AFM probes and substrates

The surface functionalization was performed as described previously^21,35,36^. Briefly, The MLCT-Bio-DC AFM probes (Bruker) and glass substrates were immersed in 50 % isopropanol for 5 mins, cleaned in a UV radiation and ozone (UV-O) cleaner and silanized with (3-aminopropyl)-triethoxysilane (APTES). A functional polyethylene glycol (PEG) cross-linker, NHS-PEG-NHS (20 kDa), was used to attach the S protein (0.1 mg/mL) and ACE2 (0.1 mg/mL) to glass substrates and AFM probes, respectively. AFM probes and glass substrates were rinse with PBS buffer (pH 7.4) for three times and immediately used for force spectroscopy experiments.

### Single-Molecule Force Spectroscopy Experiments

Atomic force microscope (Nanowizard 4, JPK) was used to acquire the force-extension curves. The experiments were performed in 10mM PBS buffer (pH 7.4). The spring constant of the cantilever was calculated by measuring its sensitivity and the thermal noise analysis, ranging from 0.04 to 0.06 N/m. The probe modified by S protein approached to substrate at a constant velocity, contacted with the ACE2 modified substrate for 0.1s, and retracted away from substrate at the same velocity. The force-extension curves recorded were analyzed using Igor 6.37 software. The unbinding rate constant and the distance from native state to transition state were estimated through Monte Carlo simulation and Bell-Evans model as described previously.^22-24,37^

### Single-Cell Force Spectroscopy

Nanowizard 4 equipped with a motorized stage (JPK Instruments) and a pump (FluidFM^®^ MFCS V2), mounted on an inverted optical microscope were used for SCFS experiments. The temperature was set to 37 °C by a Petri dish heater (JPK Instruments). The probe is a FluidFM^®^ micropipette with an aperture of 4 um and initial specified stiffness of 0.3 N/m. Each probe was calibrated before the measurement using the thermal noise analysis. To acquire a single cell to the cantilever, overnight cultivated Vero and HEK293T cells were detached from the culture flask with 0.25% (w/v) trypsin for up to 2 mins. Single cells were adsorbed to the probe by applying negative pressure of 200 mbar to the cells. When the single cell was adsorbed to the probe stably, a pressure of 50 mbar was employed to stabilize cell adsorption and avoid cell rupture caused by high pressure.

In the SCFS experiment, the cantilever attached with a single cell was approached to the S protein coated substrate at a velocity of 2 um/s. After contacting the S protein for 5 s at contact force of 2 nN, the cantilever was retracted at the same speed for 30 ums until the cells were thoroughly separated from the substrate. After one cycle of measurement, the probe with cell absorbed was immersed in 10% (v/v) sodium hypochlorite solution for 2 mins, aiming to clean the probe of any remaining cell debris, during which positive pressure of 999 mbar was applied continuously to make the probe excrete cells. The probe is then washed again in water and another fresh cell was absorbed and interacted with the substrate. Adhesion forces were quantified after the force-extension curves were baseline-corrected using JPK data process analysis software. Statistical analysis was done using GraphPad Prism 8.0.2 software.

### Antibody inhibition assays

The ACE2 and the S protein was coated to the glass substrate and the AFM probe, respectively. S protein coated probe was incubated with 100uM antibody mAb for 1h. The setpoint force was 200 pN and the probe approached the substrate at a rate of 1um/s, contacted with the substrate for 0.1s, and then retracted at the same speed. The scanning range is 5um×5um, and the sample was scanned using 32×32 pixels per line. When studying the binding probability between the S protein and the Vero cells, the set point was set as 450 pN, and the tip was approached the cell at a constant speed of 5um/s, contacted with the cell for 400ms, then retracted to 2um at the same speed. The sample was scanned using 32×32 pixels per line scanning range is 5um×5um.

### Molecular Dynamics Simulations

The RBD structures model in complex with ACE2 were taken from the Protein Data Bank (Delta PDB:7w9i^16^; Omicron PDB: 7wbl^15^). Molecular dynamics simulations were carried out using GROMACS 2018.3^38^ with the charmm36m force field^39^. The structures and force fields of Delta and Omicron in complex with ACE2 were obtained by charmm-gui^40^. The system was solvated in water box using TIP3P model. Counter ions (Na^+^ and Cl^−^) were added to the simulation box to achieve electroneutrality. After neutralizing the complexes, the systems were subjected for energy minimization by adopting the steepest descent energy minimization procedure. The temperature was maintained at 303.15 K and the pressure was maintained at 1 atmosphere. The equilibration of systems was done in the NPT ensemble for 1 ns with backbone constraints. Molecular dynamics (MD) simulation was performed for 50 ns without backbone constraints. Steered molecular dynamics (SMD) simulation was performed by harmonically restraining the position of the C-terminus of ACE2 and pulling on the C-terminus of RBD at a constant pulling speed of 10.0 Å/ns for 10 ns. Data was analyzed by VMD 1.9.3^41^ program, Pymol^42^and LigPlot^+^ 1.4.5^43^.

## Acknowledgement

We thank Prof. Xiaokang Ding and Prof. Fujian Xu for their kind help with the SCFS experiments.

## Funds

This work was supported by National Natural Science Foundation of China (11902023), National Key R&D Program of China (2021YFC2103903) and Open Project Fund (pilab1902) of the State Key Laboratory of Precision Measuring Technology and Instruments (Tianjin University).

