## Supporting for "Pathogen-Host Adhesion between SARS-CoV-2 Spike Proteins from Different Variants and Human ACE2 Probed at Single-Molecule and Single-Cell Levels"

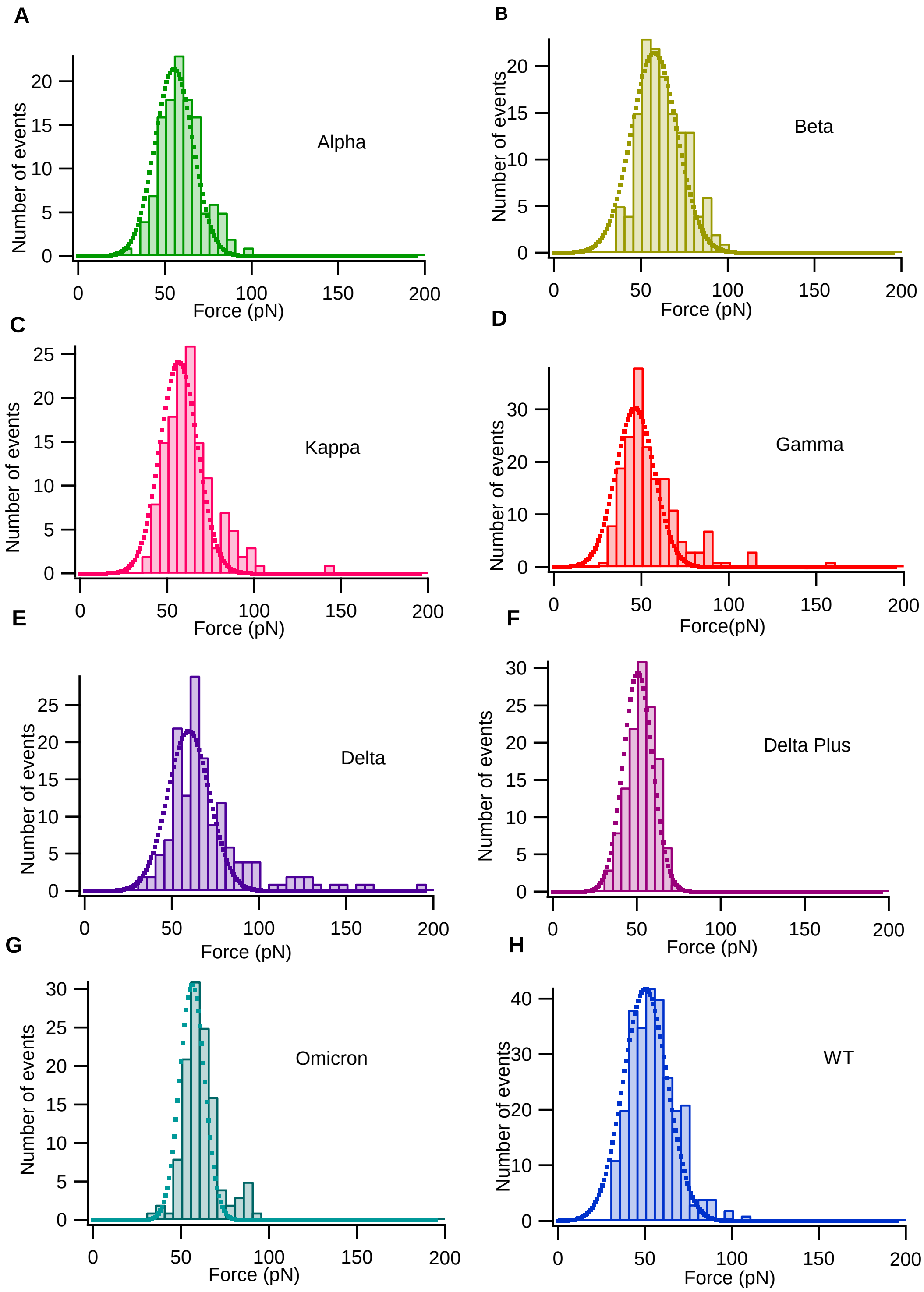


**Figure S1.** Representative histograms of rupture force distribution for the interaction between ACE2 and different S proteins. Dashed lines represent the results of gaussian fitting.

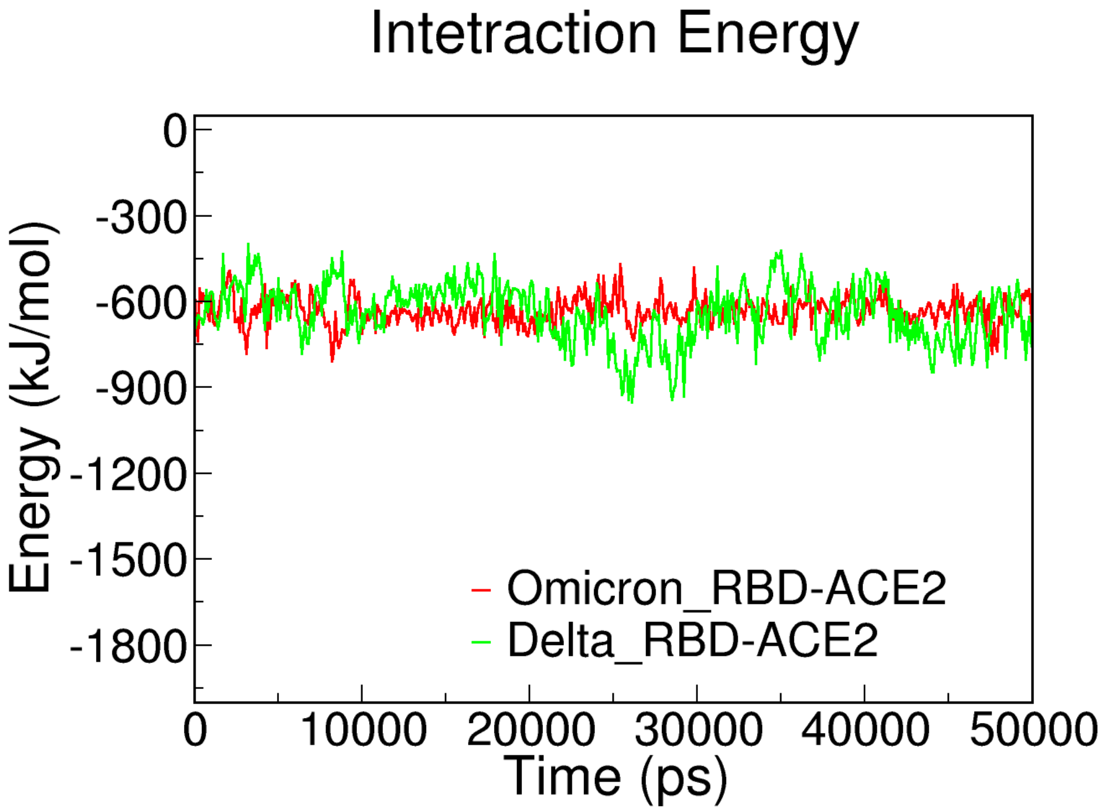


**Figure S2.** Energy diagrams in molecular dynamics simulations.

**Table S1.** Mutated amino acids in different variants of spike proteins

| Spike protein | Mutations |
|---|---|
| Omicron | A67V, HV69-70del, G142D, VYY143-145del, N211del, L212I, ins214EPE, G339D, S373P, S375F, K417N, G446S, S477N, T478K, E484A, Q493R, G496S, Q498R, N501Y, Y505H, T547K, D614G, H655Y, N679K, P681H, N764K, D796Y, N856K, Q954H, N969K, L981F |
| Delta | T19R, G142D, EF156-157del, R158G, L452R, T478K, D614G, P681R, D950N |
| Delta Plus | T19R, V70F, FR157-158Del, A222V, W258L, K417N, L452R, T478K, D614G, P681R, D950N |
| Alpha | V69-70del, Y144del, N501Y, A570D, D614G, P681H, T716I, S982A, D1118H |
| Gamma | L18F, T20N, P26S, D138Y, R190S, K417T, E484K, N501Y, D614G, H655Y, T1027I, V1176F |
| Kappa | T95I, G142D, E154K, L452R, E484Q, D614G, P681R |
| Beta | L18F, D80A, D215G, LAL242-244del, R246I, K417N, E484K, N501Y, D614G, A701V |
